# Mapping the Brazilian Scientific Diaspora: Migration Patterns of PhDs in Global Mobility

**DOI:** 10.1101/2024.04.22.590567

**Authors:** Concepta McManus, Brenno A. D. Neto, Abilio Afonso Baeta Neves, Rafael Tavares Schleicher, Claudia Figueiredo

**Author notes:** Corresponding author: Concepta McManus and Claudia Pinto Figueiredo.

## Abstract

A scientific diaspora refers to a community of scientists who have emigrated from their home country to work in another nation. This study investigates this phenome in depth the Brazilian context, examining who comprises this diaspora (doctorates, postdocs, lecturers), where they have migrated, and their areas of study. We conducted this examination based on publications by Brazilian doctors who graduated between 2005 and 2021, as well as post-doctorates and students with full doctorate scholarships abroad. These students were identified on the CAPES open data website. The publications of these authors were captured in Scopus and Web of Science. Then those with addresses abroad were analysed in Vosviewer® and using logistic regression (stayed abroad or not), area of knowledge and a decision tree to see the effect of the Brazilian university region, type of institution and scholarship on the decision to migrate. The level of diaspora is approximately 1.7% among all doctorates trained in Brazil, reaching 6.6% in postdoctoral scholars with experience abroad. This suggests that PhDs with advanced training and experience have a higher propensity to emigrate from Brazil. These PhDs predominantly choose to migrate to North America and Western Europe, with a strong preference for careers in Science, Technology, Engineering, and Mathematics (STEM) fields. Brazilian PhDs with international experience tend to have a more diverse migration pattern, while those who complete their PhD in Brazil show a distinct preference for migrating to Portugal, indicating differing global mobility based on scientific experience. A decision tree analysis reveals that life or exact sciences PhDs, those who graduated after 2012, and obtained their postgraduate degrees from institutions in the southeast or south of Brazil are more likely to migrate. While the reasons behind these migration patterns are not evaluated in this study, better job prospects, higher salaries, or more substantial research funding could be influential factors in the decision to migrate.

## Introduction

Diaspora is a Greek word that means to sow or scatter. As applied to people, the Greek historian Thucydides probably first used the term to describe the dispersion of the Greeks. Today, it basically refers to the dispersion or scattering of a people with common ethnic or cultural origins who find themselves living in different geographic locations. The high international mobility of academics in recent decades is becoming a key feature of the global scientific community. Many successful scientists were born in one country, educated in another, conducted research in a third country, and taught in a fourth. In some scientific areas, participation in international academic mobility (even temporary jobs or internships abroad) is almost an obligatory condition for a successful scientific career.

The impact of scientific diaspora on a country’s scientific landscape is nuanced, involving both positive and negative elements (Song, 2014). “Brain Gain” is a key advantage, as skilled scientists returning to their home country contribute valuable knowledge, fostering scientific community growth. Furthermore, it significantly fuels innovation by introducing diverse perspectives, methodologies, and approaches. The networking capabilities of diaspora scientists create global connections, promoting collaboration and resource sharing. Lastly, the involvement of diaspora scientists in obtaining funding for scientific research can significantly benefit their home country. Their international networks and expertise enhance the country’s competitiveness in securing financial support for research. The drawbacks of scientific diaspora include “Brain Drain,” where the departure of talented scientists depletes essential talent and resources in the home country. Disconnection arises as diaspora scientists may become less engaged in local scientific developments, hindering knowledge exchange. Loss of local expertise occurs when scientists leave specific regions, impacting overall scientific capacity. Competition arises as diaspora scientists may be perceived as competitors, reducing collaboration. Additionally, the global contributions of diaspora members may not always benefit their home countries, posing challenges in maximizing positive impacts. Effectively managing these dynamics is crucial for cultivating a sustainable and mutually beneficial scientific environment.

The Brazilian diaspora in science refers to Brazilian scientists and researchers who have left the country for various reasons, including seeking better professional opportunities, lack of funding and resources in Brazil, and political or economic instability. Despite the challenges, many Brazilian scientists abroad have made significant contributions in their fields of study, such as medicine, engineering and natural sciences. Some researchers live and work in two countries. Some emigrant scientists later return to their homeland. Others stay there forever. As it is often difficult to distinguish between these groups, state measures to interact with them must be interrelated. However, there is little information about Brazil’s scientific diaspora in quantitative terms, areas of knowledge and countries of work. As a rule, the diffusion of knowledge is traced through citations of patents and scientific publications (Yurevich et al., 2019).

In this study, we tried to identify the Brazilian scientific diaspora by identifying Brazilian PhDs with affiliation only abroad in various international publishing platforms. The questions asked include i) how many alumni move abroad; ii) what factors including scholarship type, areas of knowledge, type of original and host institution and are more likely to induce moving abroad.

## Results

A total of 268,847 PhDs trained in Brazil in the period, with 143,809 scholarship holders abroad (all scholarships including masters and undergraduate), 97,052 university lecturers and 42,588 researchers with addresses in Brazil and abroad were identified, with a total of 437,987 individual identities (Table 1). We identified 4,631 researchers that receiving a doctorate in Brazil and address abroad (1.72% of the doctors trained in Brazil in the period). Table 1 summarizes some important information. The overall level of diaspora is 1.66% of the doctors trained in the period (2005-2021).

**Table 1.** Profile of Brazilians PhD that defending doctorates between 2005 and 2021, including information on Scholarship Abroad, Postgraduate Supervisor Roles in Brazil, address linked to Scientific Publications.

| PhD in Brazil | Received Scholarship Abroad | Postgraduate Supervisor in Brazil | Address Abroad in Publications | Total |
| --- | --- | --- | --- | --- |
| x | x | x | x | 1090 |
| x | x | x |  | 9872 |
| x |  | x | x | 1390 |
| x |  | x |  | 40102 |
| x | x |  |  | 27393 |
|  | x | x | x | 993 |
|  | x | x |  | 7611 |
|  |  | x | x | 2229 |
|  |  | x |  | 33691 |
| x |  |  |  | 183345 |
|  | x |  |  | 93611 |
|  |  |  | x | 31129 |
|  | x |  | x | 886 |
| x | x |  | x | 1706 |
| x |  |  | x | 2925 |

Source: Capes, CNPq, Fapesp and Web of Science

There were 267,823 Brazilian doctors who graduated with PhD from 2005 to 2021. Of these, 2925 were identified as living abroad without having a period abroad during their studies (Fig. 1a), while 1706 did have a scholarship abroad (Fig. 1b). These were identified as having studied mainly in North America and Western European Countries (Figure 1b). The countries where Brazilians (33,300) studied for up to a year abroad during their doctorate (sandwich doctorate) period are shown in Figure 1c. The central countries are in the global north, especially in the United States and Western Europe. 886 students received scholarships for full PhD abroad. The present study identified 7608 scholarship holders with a full doctorate or post-doctorate abroad without affiliation in Brazil.

**Figure 1.**
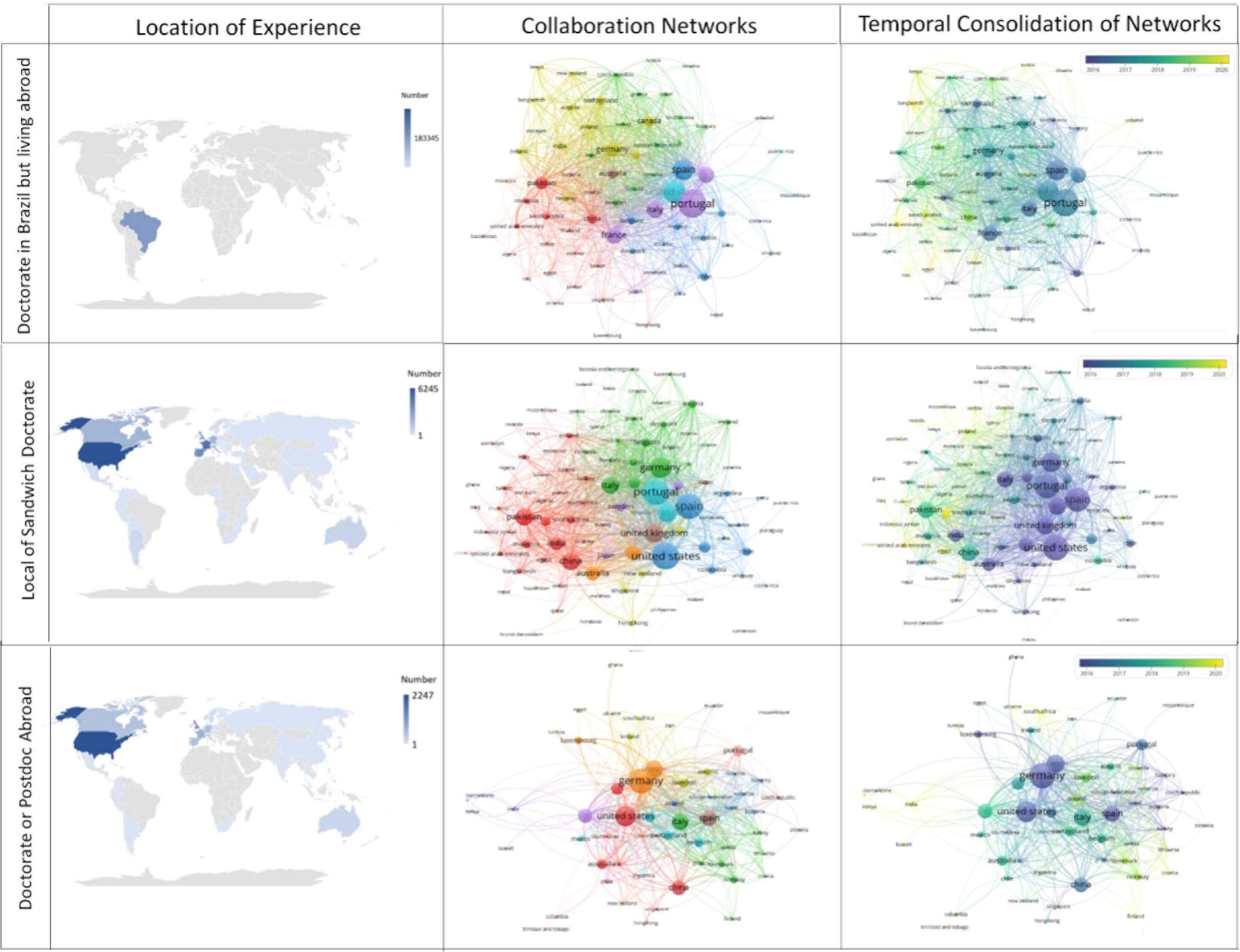
Collaboration Networks of Brazilian PhD Graduates that defending doctorates between 2005 and 2021 with International Affiliations. The left column maps the geographical experience of these researchers, with the top map highlighting those whose experience was confined to Brazil, the middle map showing countries where researchers engaged in ‘sandwich’ doctoral programs, and the bottom map indicating countries where researchers completed their full doctoral or postdoctoral studies. The central column reveals the current scientific collaboration networks of these Brazilian PhDs, while the right column displays the temporal depth of these networks. The colour coding in the networks ranges from yellow for recent collaborations to blue for longer-standing ones, illustrating the growth and durability of international scholarly connections.

The networks of countries linked to these researchers are coded by clusters with each colour indicates a cluster of countries (Fig. 1, left column). As above, there is a concentration in Western European countries such as Portugal, Spain and the United States. Those that carried out their PhD in Brazil but without time abroad show a concentration in the Iberian Peninsula, Western Europe and North America.

The primary affiliation institutions for Brazilians abroad show the importance of Portuguese and French Higher Education Institutions (HEIs). In the right column of Figure 1 illustrates countries with more recent affiliations (Yellow colour), like Israel, China and Saudi Arabia, which tend to show more citations. Full doctorates and postdocs have the highest affiliation in Germany, France, and the US, which are older affiliations. Among the top Higher Education and Research Institutions, a variety of institutions from different countries, including the US, France, Germany, and others, are represented. There are former students in private companies (Xerox) and research centers (CNRS and Helmholtz), as well as universities. The full doctoral and postdoctoral affiliation research groups show few links between the groups, indicating that deciding to stay abroad is not necessarily about building diaspora networks.

Regarding length of affiliation, the same countries are indicated as having the oldest affiliation. Affiliation with Asian countries is more recent (Figure 1, right column), and in general, Brazilians are not group leaders. There are groups for medicine (light blue), physics (green), computer science (yellow), chemistry (dark blue), and engineering (red), among others (Figure 1. left column). There are more recent groups in the areas of chemistry and physics, as well as areas of basic science.

Looking at the 4,631 former Brazilian PhD students, there was a constant increase until 2018, with stabilization after this date. The southeast and southern regions (Figure 2b) produce more former students abroad, especially in São Paulo, Rio Grande do Sul, Minas Gerais and Rio de Janeiro (Figure 2c). The three state universities in São Paulo, in addition to UFRGS, train more students abroad (Annex 2). There is a greater proportion of former students from programs with higher grades (Figure 2d) with addresses abroad, with 75% with grades 5 or above.

**Figure 2.**
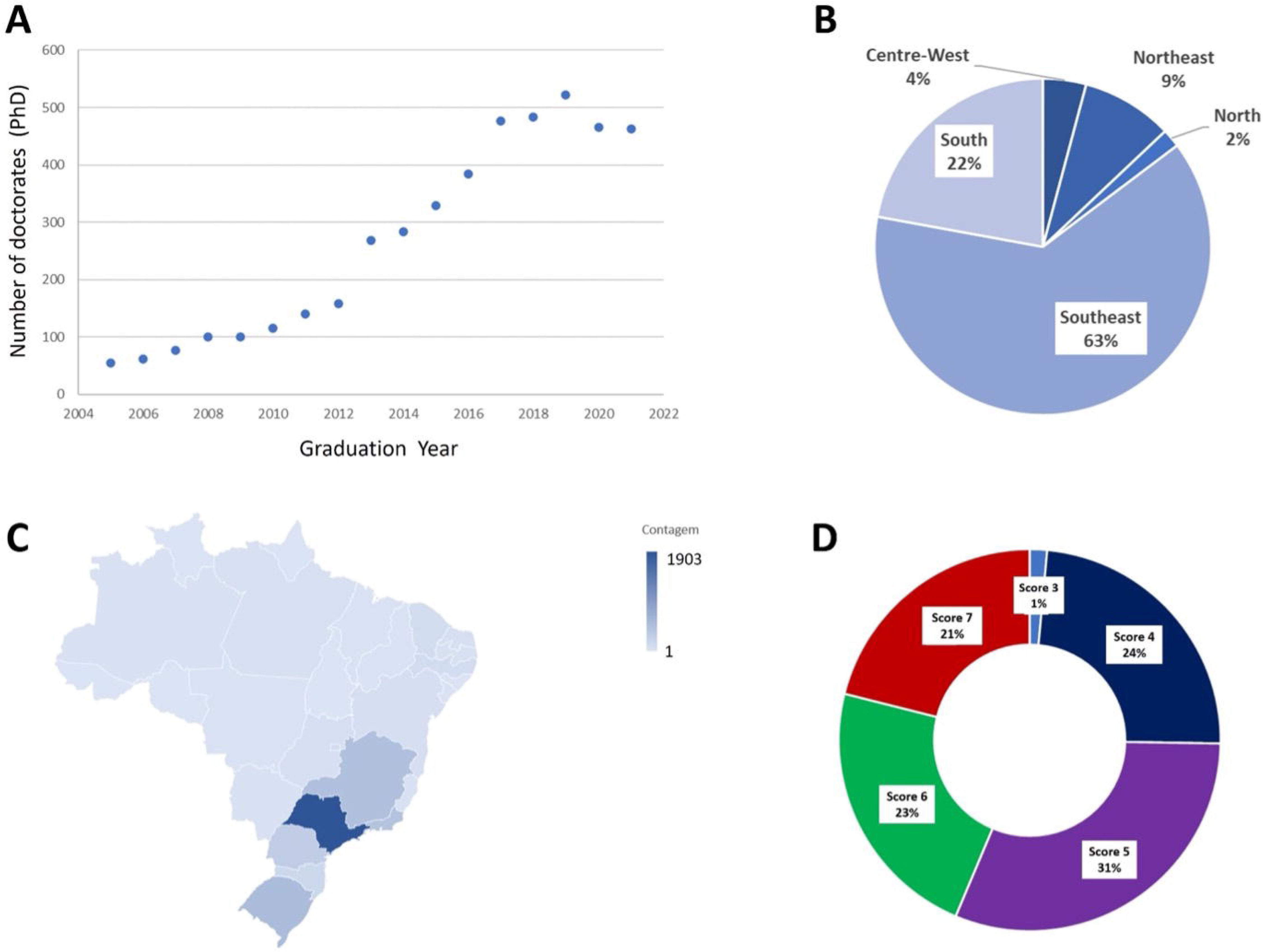
Profile of Brazilian PhD Graduates that defending doctorates between 2005 and 2021 with International Affiliations. A) The number of Brazilian PhD holders who have published in the Web of Science from 2004 to 2021. B) Percentage distribution of these PhD holders by Brazilian regions, highlighting the Southeast’s predominant 63%. C) Statewide distribution within Brazil. D) CAPES evaluation scores for Brazilian graduate programs, ranging from 3 to 7, with 7 being the highest mark for quality and 3 the lowest. Data sources: Scopus, Web of Science, CNPq, CAPES, and FAPESP.

The distribution of research specializations among former Brazilian PhD students demonstrates a significant inclination towards the Life Sciences, which encompasses the majority with 64% representation. This is followed by the Exact Sciences, accounting for 29% of the areas of study. Social Sciences and Humanities are studied to a lesser extent, making up 7% of the research focus among these PhD graduates. Artificial intelligence (AI) and promising machine learning (ML) techniques from computer science are widely affecting many aspects of various fields, including science and technology, industry and even our everyday lives (Figure 3). This data points to a strong emphasis on life and exact sciences within the Brazilian PhD community.

**Figure 3.**
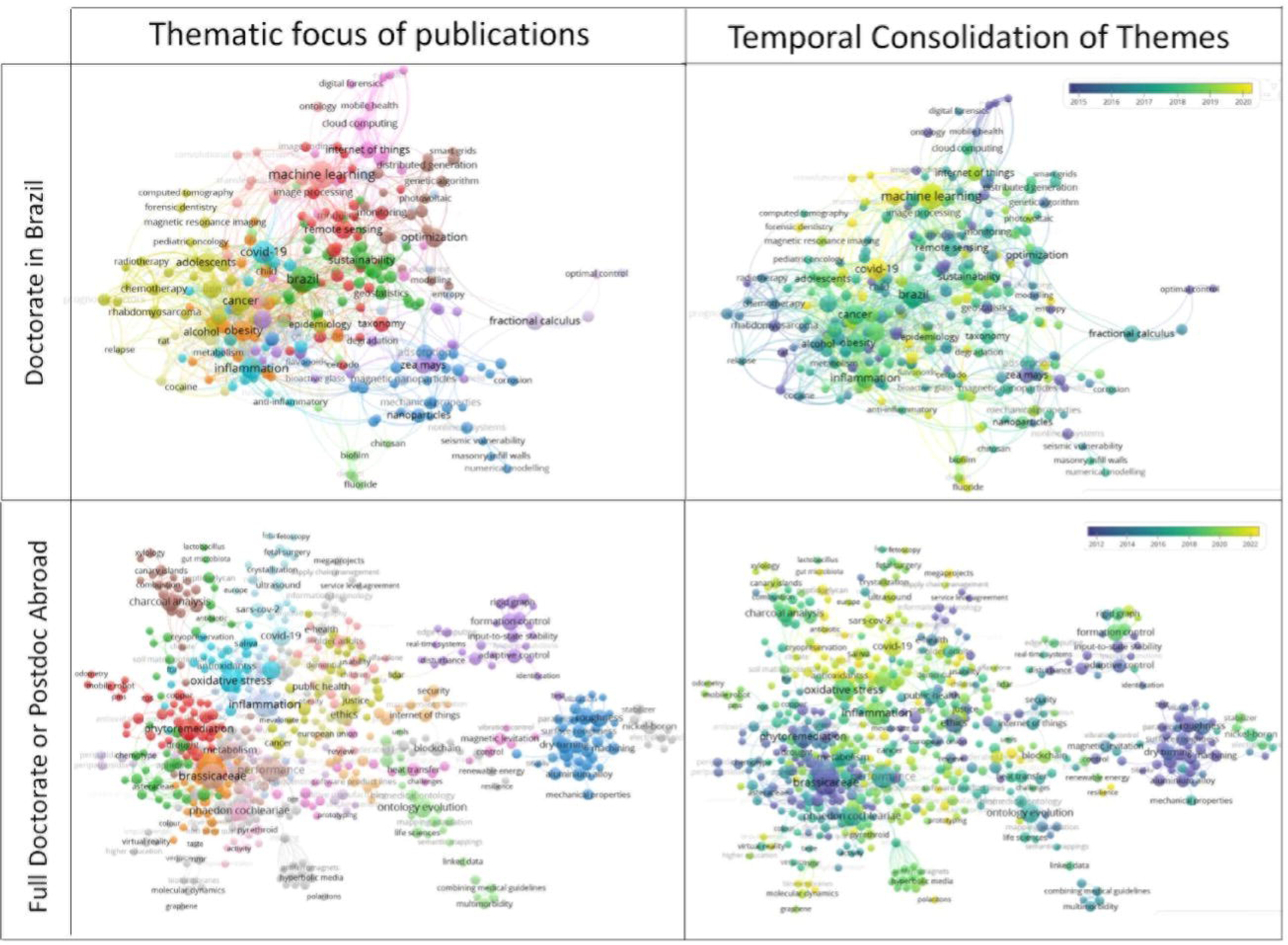
Thematic Distribution of International Publications by Brazilian PhD Graduates (2005-2021). The top panels represent the thematic focus of publications by Brazilian PhD holders who completed their degrees in Brazil and have since published with International Affiliations. The bottom panels detail the themes of publications by Brazilian PhD holders who completed full doctoral studies or postdoctoral research abroad. The right column displays the temporal depth of the thematic focus and the color coding ranges from yellow for recent collaborations to blue for longer-standing ones, illustrating the temporal consolidation of knowledge areas. Data were sourced from the Scopus database.

The areas of knowledge of former and post-doctorates are in exact areas such as engineering and computer science, as well as in areas of life such as medicine and biochemistry. This is per NSF (2012) in the USA.

The areas of knowledge of former Brazilian PhD students (Figure 3) show concentration in Life followed by exact sciences. The themes studied (Figure 4) show this trend. Artificial intelligence (AI) and promising machine learning (ML) techniques from computer science are widely affecting many aspects of various fields, including science and technology, industry and even our everyday lives.

**Figure 4.**
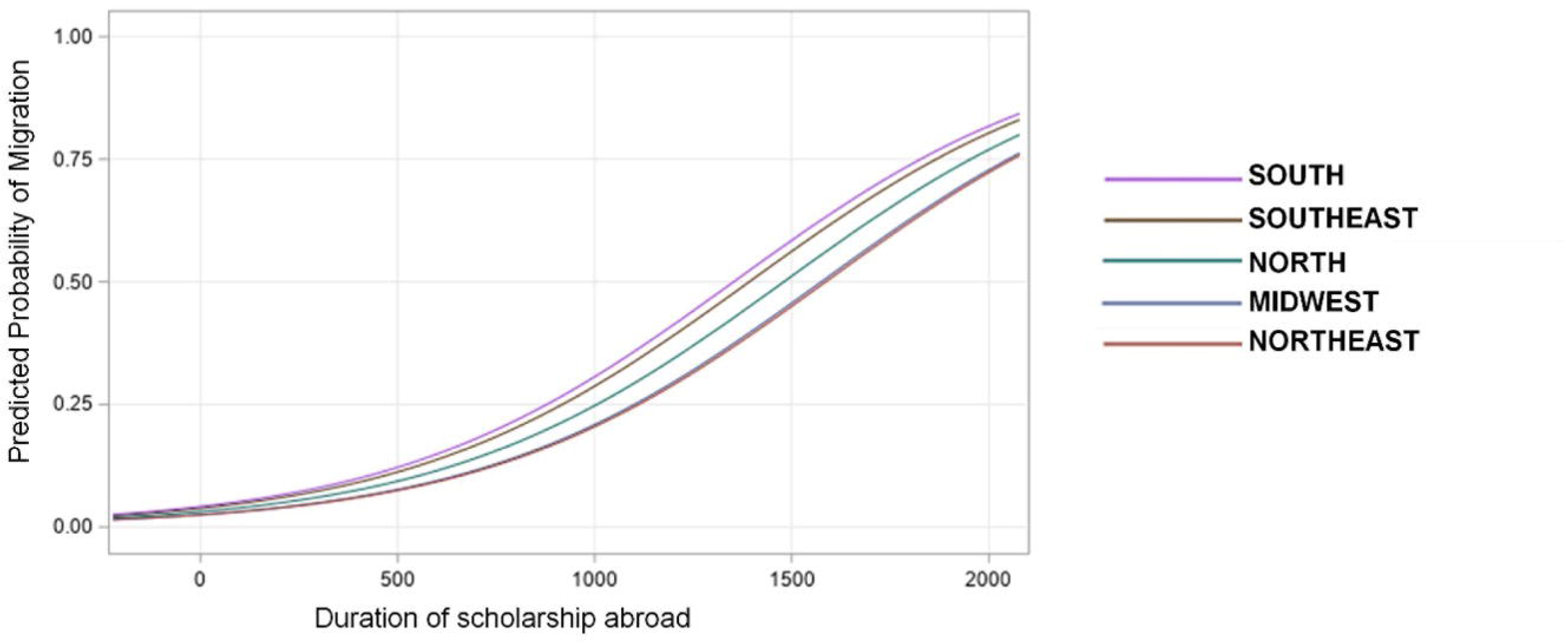
Migration Probability Curve for Brazilian Scholars. The graph demonstrates the predicted probability of migration, illustrating the likelihood of Brazilian PhD migrating based on the time abroad and the regional location of their graduate program. This probabilistic model is derived from data across multiple scholarly databases including Scopus, Web of Science, CNPq, Capes, and FAPESP.

For logistic regression, the ROC curve has an area under the curve of 0.70, showing the adequacy of the analysis model. There is a significant effect (P<0.01) of the graduation year (OR = 1.09; CI = 1.08 to 1.10) and program score (OR = 1.25, CI = 1.21 to 1.29) on the possibility of going abroad with a student who graduated in more recent years and higher concept having higher probability. Table 2 shows differences between the country’s regions, area of knowledge and type of HEI for the probability of going abroad. Former students from the Northeast and North regions are less likely to stay abroad, as well as those from programs with a grade lower than 5 and former students from private HEIs.

**Table 2.** Odds ratios for the probability of PhD migrating.^1^.

| Region of HEI of Doctorate |  |  |  |  |  |
| --- | --- | --- | --- | --- | --- |
|  | CO | NE | N | SE |  |
| NE | 1.26** |  |  |  |  |
| N | 1.07 | 0.85 |  |  |  |
| SE | 0.71** | 0.56** | 0.66** |  |  |
| S | 0.61** | 0.49** | 0.57** | 0.86** |  |
| Judicial Status of The Institution |  |  |  |  |  |
|  | State | Federal | Municipal |  |  |
| Federal | 1.15* |  |  |  |  |
| Municipal | 2.56 | 2.22 |  |  |  |
| Private | 2.50** | 2.17** | 0.98 |  |  |
| Knowledge Area |  |  |  |  |  |
|  | Exact | SSH |  |  |  |
| SSH | 4.52** |  |  |  |  |
| Life | 0.76** | 0.16** |  |  |  |
| Region |  |  |  |  |  |
|  | Africa | North America | South America | Asia | Europe |
| N. America | 0.53 |  |  |  |  |
| S. America | 1.25 | 2.37** |  |  |  |
| Asia | 0.83 | 1.56 | 0.66 |  |  |
| Europe | 0.68 | 1.28* | 0.54** | 0.82 |  |
| Oceania | 0.43** | 0.81 | 0.34** | 0.52* | 0.63** |
| Scholarship Type Abroad |  |  |  |  |  |
|  | Senior Scientist | Full Doctorate | Post-doc |  |  |
| Full Doct | 1.15 |  |  |  |  |
| Post doc | 1.34* | 0.99 |  |  |  |
| Sandwich | 2.01** | 1.74** | 1.77** |  |  |

\*P<0.05; \*\*P<0.01; HEI – Higher Education Institute, SSH – Social Sciences and Humanities.
<sup>1</sup>An odds ratio greater than one means the horizontal variable is more likely to occur than the vertical variable.

Time and the doctoral region (Southeast and South) increase the possibility of staying abroad (Figure 4). The number of articles published by former students abroad (Figure 5) shows a constant growth since 2005. The location of Brazil’s full doctorate or postdoctoral fellows abroad (Table 3) shows the preference for the US and Germany, among other Western European countries.

**Figure 5.**
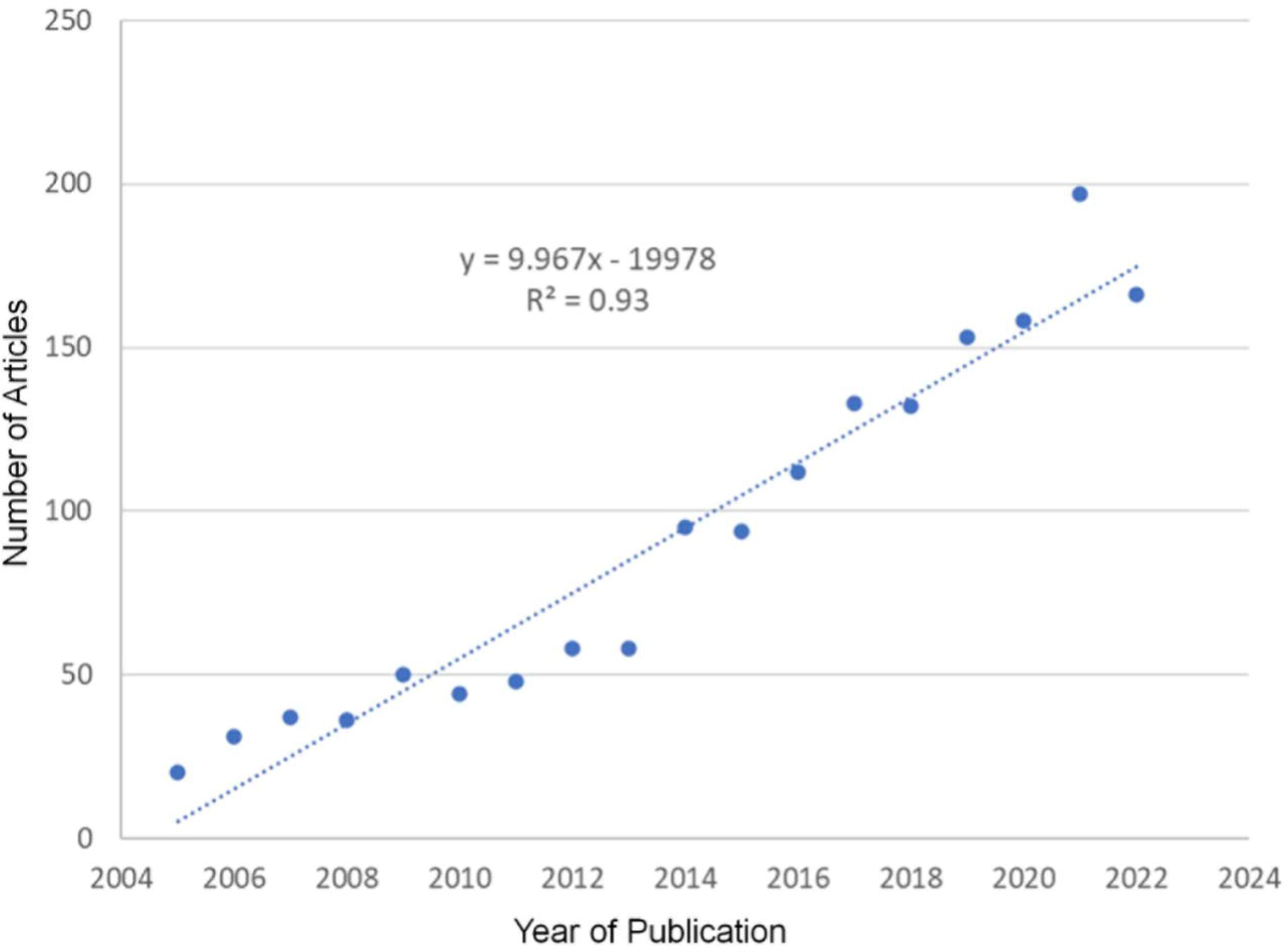
Trends in Scholarly Publishing. This figure illustrates the publication trajectory over time, measuring the annual output of articles published by former Brazilian PhD students with international affiliations. The dataset is divided into two categories: (a) those who participated in ‘Sandwich’ PhD programs, and (b) those who completed full PhDs or conducted postdoctoral research abroad. The fitted line provides a regression model indicating the growth trend, with the equation and R² value denoting the strength of this trend over the specified period.

**Table 3.** Knowledge area and funders of studies of Brazilian PhD alumni.

| Full or PD |  | Sandwich |  | Full or PD |  | Sandwich |  |
| --- | --- | --- | --- | --- | --- | --- | --- |
| Country | Docs | Country | Docs | Subject area | Docs | Subject area | Docs |
| Germany | 449 | Portugal | 1,591 | Computer Science | 905 | Engineering | 2603 |
| United States | 396 | United States | 970 | Engineering | 873 | Computer Science | 2381 |
| France | 217 | Spain | 839 | Medicine | 635 | Medicine | 1969 |
| Italy | 187 | France | 542 | Mathematics | 477 | Agricultural and Biological Sciences | 1701 |
| China | 170 | Italy | 530 | Biochemistry, Genet. & Mol. Biology | 403 | Biochemistry, Genetics and Molecular Biology | 1197 |
| Spain | 167 | United Kingdom | 494 | Physics and Astronomy | 335 | Mathematics | 1182 |
| United Kingdom | 163 | Germany | 432 | Chemistry | 309 | Physics and Astronomy | 1020 |
| Canada | 106 | Pakistan | 297 | Environmental Science | 293 | Environmental Science | 978 |
| Portugal | 84 | Canada | 231 | Social Sciences | 290 | Social Sciences | 879 |
| Netherlands | 69 | China | 178 | Materials Science | 272 | Chemistry | 850 |
| Australia | 65 | Australia | 171 | Agricultural & Biological Sciences | 231 | Materials Science | 811 |
| Switzerland | 64 | Switzerland | 165 | Energy | 203 | Energy | 633 |
| Belgium | 61 | Netherlands | 132 | Chemical Engineering | 172 | Earth and Planetary Sciences | 611 |
| Sweden | 61 | Chile | 129 | Earth and Planetary Sciences | 151 | Chemical Engineering | 507 |
| Norway | 37 | Belgium | 117 | Pharmac., Toxicology & Pharmaceuticals | 135 | Pharmacology, Toxicology and Pharmaceuticals | 432 |
| Luxembourg | 36 | Argentina | 101 | Decision Sciences | 92 | Immunology and Microbiology | 359 |
| Austria | 33 | Mexico | 85 | Arts and Humanities | 91 | Neuroscience | 277 |
| Denmark | 30 | Saudi Arabia | 82 | Business, Management & Accounting | 91 | Business, Management and Accounting | 275 |
| Ireland | 28 | India | 81 | Neuroscience | 89 | Decision Sciences | 271 |
| Mexico | 26 | Sweden | 81 | Immunology & Microbiology | 76 | Veterinary | 210 |
| Poland | 19 | Norway | 78 | Economics, Econometrics & Finance | 69 | Arts and Humanities | 202 |
| Japan | 17 | Colombia | 77 | Nursing | 51 | Nursing | 186 |
| Chile | 15 | Austria | 75 | Multidisciplinary | 45 | Economics, Econometrics and Finance | 153 |
| Greece | 15 | Japan | 69 | Health Professions | 42 | Multidisciplinary | 146 |
| Czech Republic | 12 | Malaysia | 64 | Psychology | 36 | Dentistry | 143 |
| South Africa | 12 | Denmark | 63 | Veterinary | 17 | Health Professions | 142 |
| Colombia | 9 | Poland | 55 | Dentistry | 8 | Psychology | 114 |
| Israel | 9 |  |  |  |  |  |  |
| Turkey | 9 |  |  |  |  |  |  |

The result of the decision tree (Figure 6a) showed that the critical factors for migrating abroad were i) being from the area of life sciences or exact sciences, ii) having graduated after 2012, and iii) having a postgraduate degree at a university in the southeast or south region.

**Figure 6.**
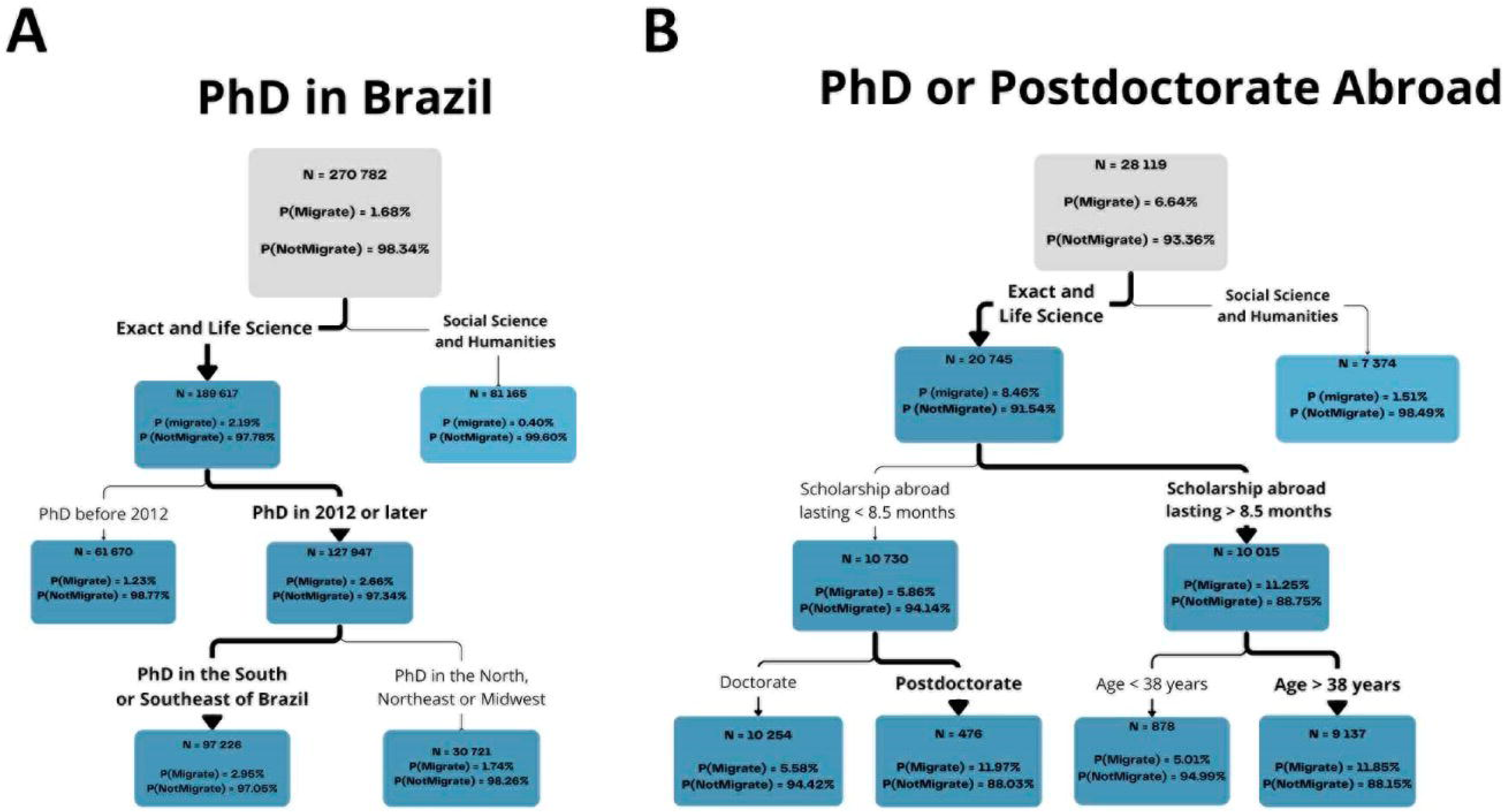
Decision Tree for the Academic Migration of Brazilian PhD Graduates (2005-2021). Each node (box) provides the sample size (N) and the probability percentages (P) for migration and non-migration outcomes. **(A)** Decision tree tracing the academic migration patterns of Brazilian PhD holders who completed their degrees in Brazil (N=270,782). Data were segmented by critical factors such as field of study (Exact and Life Science), date of degree completion (PhD before 2012 or PhD in 2012 or later), geographic origin of post-graduation program Brazil (South or Southeast of Brazil or North, Northeast, or Midwest of Brazil. **(B)** Decision tree tracing the academic migration patterns of Brazilian PhD holders who completed full doctoral studies or postdoctoral research abroad (N = 28,119). Data were segmented by critical factors such as field of study (Exact and Life Science), duration of Scholarship Abroad (lasting < 8.5 months or lasting ≥ 8.5 months), type of scholarship (doctorate or postdoctorate), age at time of experience abroad (<38 years or ≥38 years).

The decision tree (Figure 6b) for doctorates and postdocs shows that the area of knowledge, time spent abroad, type of scholarship and age of the candidate significantly affect the decision to migrate. In this case, there is a higher level of migration (around 6.6%). It is worth remembering that this is the percentage of post-doctors with experience abroad, not the total level of post-doctors, as this information was not available. Once again, being higher in the exact and life areas, especially after a more extended scholarship abroad (>8.5 months), senior internship type scholarship and with more advanced age (>38 years). In part, the reasons may include a lack of prospects for public university employment (rising retirement age means current professors are staying longer (McManus et al., 2023), lack of research resources, or parents of young families wanting “a better life”, among others.

The United Nations’ Sustainable Development Goals (SDGs) represent a universal call to action to end poverty, safeguard the environment, and secure a prosperous and equitable future for all by 2030. Embedded in these 17 goals is an integrated approach that recognizes the interplay between social equity, economic growth, and environmental protection. The SDGs ambitiously target a variety of critical global challenges, including but not limited to poverty, hunger, health disparities, and gender inequality. Achieving these goals necessitates widespread collaboration and resources from various societal sectors. An examination of the scholarly contributions by Brazilian PhDs with affiliation abroad offers insightful reflections on the alignment of academic themes with the SDGs. Our analysis reveals a prominent focus on Good Health and Well-being (SDG 3), which emerges as the predominant theme among the research outputs of the Brazilian academic diaspora (Figure 7a), followed by climate action and life on earth.

**Figure 7.**
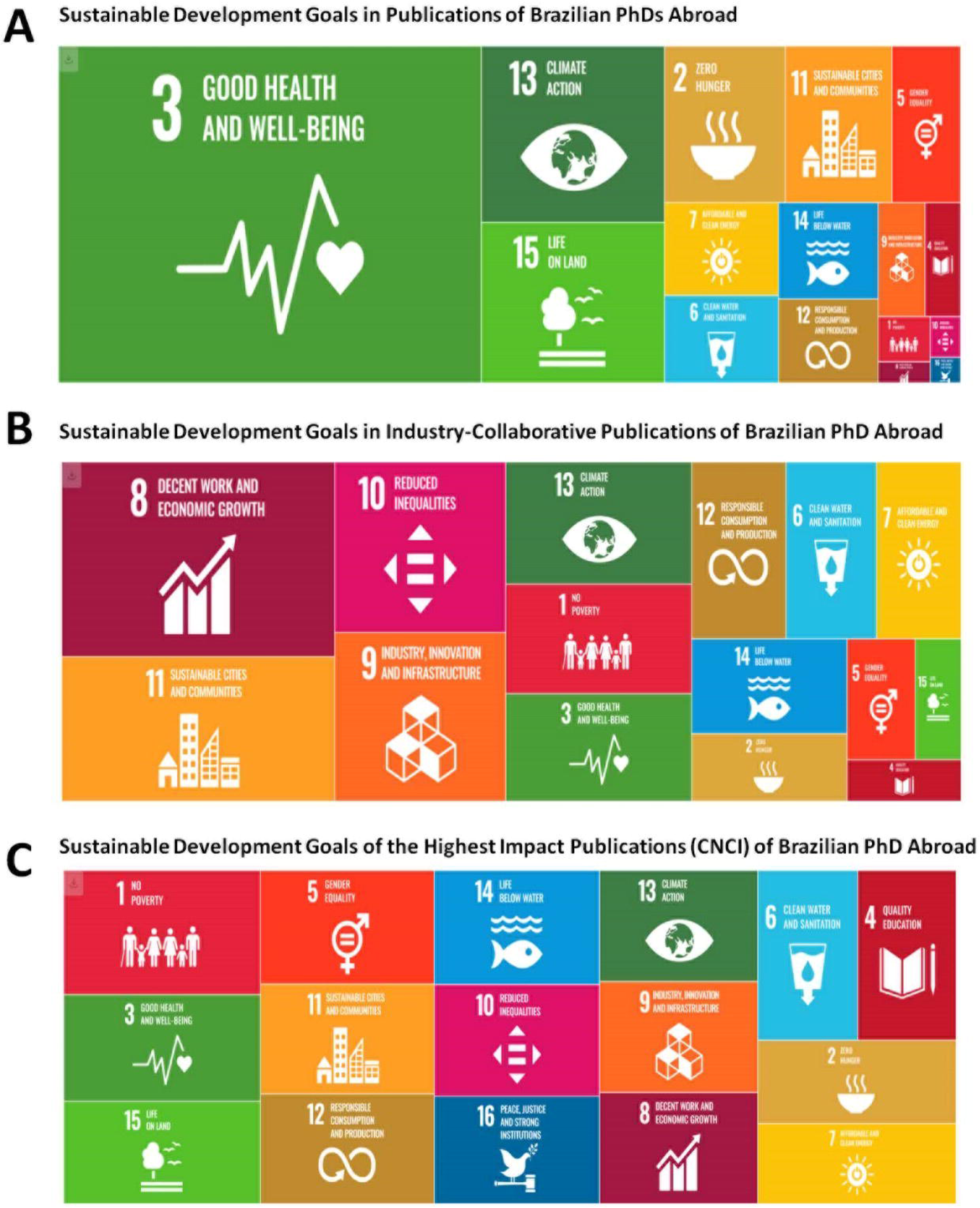
Analysis of publications by Brazilian PhD abroad, categorized by the Sustainable Development Goals (SDGs) adopted by the United Nations. Each SDG is represented by a distinct colour and icon, with corresponding numerical indicators for each of the three analysis parts, offering a comprehensive view of the contribution and impact of Brazilian researchers in the international context. **(A)** Total number of papers contributed to each SDG, providing insights into research volume and focus areas in Publications of Brazilian PhDs Abroad (2005-2021). **(B)** Proportion of these papers that were co-authored with industry partners, indicating the extent of academia-industry collaboration within each SDG.SDG in Industry-Collaborative Publications of Brazilian PhD Abroad. **(C)** SDG of the Highest Impact Publications (CNCI) of Brazilian PhD Abroad. Data were sourced from Scopus, Web of Science and Capes.

Industry collaborations, which are instrumental in translating academic research into practical solutions, show a pronounced orientation towards Decent Work and Economic Growth (SDG 8) and Sustainable Cities and Communities (SDG 11) (Figure 7b). When assessing the impact of these publications using the Category Normalized Citation Impact (CNCI), a notable trend emerges. Publications pertaining to No Poverty (SDG 1), Health (SDG 3), and Life on Land (SDG 15) demonstrate the highest impact (see Figure 7c). The significant CNCI associated with these SDGs illustrates the potent role that research can play in informing and catalysing actions aimed at eradicating poverty, enhancing global health, and protecting biodiversity. Collaborations with industry (Figure 7b) were highest in the areas of work (SDG8) and sustainable cities (SDG11), highlighting the potential of industry to contribute to economic development and urban sustainability. When assessing the impact of these publications using the Category Normalized Citation Impact (CNCI), a notable trend emerges. Publications pertaining to No Poverty (SDG 1), Health (SDG 3), and Life on Land (SDG 15) demonstrate the highest impact (see Figure 7c).

Table 4 presents a comparative analysis of the research impact of Brazilian PhDs residing in Brazil versus those abroad, with metrics aligned to the Sustainable Development Goals (SDGs). The color-coding system—blue for PhDs living in Brazil and pink for those abroad—illustrates relative performance across several indicators. Brazilian PhDs abroad show stronger performance in being cited and having a higher world relative impact. Brazilian PhDs with affiliation in Brazil show stronger performance in the number of documents cited with a higher percentage of open access publications. Comparing the publications from the diaspora with Brazil in general (Table 4), we can observe a worsening in the indicators, suggest that Brazil has been lose quality researchers abroad in themes of relevance for the country’s development and fulfilment of international agreements. However, this comparison has significant limitations, as it contrasts the performance of PhDs within highly disparate realities concerning funding and working conditions. This trend (Figure 7 and Table 4) has implications for the ability of Brazil to contribute to the SDGs, but demonstrated that Brazilian PhDs abroad, especially in collaboration with industry, can drive progress in sustainable development.

**Table 4.** Comparative Analysis of Research Impact Among Brazilian PhDs abroad: A Study Based on Sustainable Development Goals (SDGs)

| Differences | % Docs |  | % Docs |  | % Corresp. Author | % |  | % Gold Docs | % Docs Q1 |
| --- | --- | --- | --- | --- | --- | --- | --- | --- | --- |
|  | Cited | CNCI | Top 10% | % Ind. Collab |  | Highly Cited | IRW |  |  |
| 01 No Poverty | -8.80 | -47.51 | -61.00 | 4.08 | 37.05 | -91.35 | -59.85 | 14.42 | -35.64 |
| 02 Zero Hunger | -3.57 | -74.51 | -139.78 | -144.32 | 34.90 | -222.22 | -78.43 | 42.00 | -90.87 |
| 03 Good Health and Well-being | -3.64 | -59.83 | -98.89 | -187.68 | 39.76 | -160.00 | -60.12 | 21.86 | -48.05 |
| 04 Quality Education | -14.86 | -57.51 | -88.74 | -145.45 | 38.61 | -130.00 | -64.20 | 21.79 | -50.54 |
| 05 Gender Equality | -4.69 | -58.44 | -93.85 | -196.30 | 38.05 | -162.50 | -59.92 | 27.73 | -48.70 |
| 06 Clean Water and Sanitation | -2.86 | -32.89 | -58.12 | -15.50 | 35.71 | -100.00 | -30.22 | 27.34 | -33.41 |
| 07 Affordable and Clean Energy | -7.39 | -28.41 | -49.34 | -56.52 | 36.99 | -86.67 | -32.45 | 5.74 | -19.38 |
| 08 Decent Work and Economic Growth | -8.14 | -25.96 | -50.29 | -10.71 | 39.69 | -104.35 | -42.56 | 25.19 | -33.31 |
| 09 Industry, Innovation and Infrastructure | -9.51 | -34.02 | -47.69 | -106.05 | 34.58 | -76.74 | -46.46 | -1.19 | -19.36 |
| 10 Reduced Inequality | -13.11 | -42.40 | -75.53 | 5.99 | 40.17 | -166.67 | -47.70 | 34.57 | -25.52 |
| 11 Sustainable Cities and Communities | -6.54 | -38.35 | -54.50 | -78.39 | 37.73 | -97.87 | -44.50 | 9.06 | -23.90 |
| 12 Responsible Consumption and Production | -4.07 | -32.58 | -60.33 | -80.00 | 34.34 | -135.71 | -36.53 | 20.43 | -31.51 |
| 13 Climate Action | -3.89 | -56.22 | -104.14 | -82.00 | 35.72 | -141.30 | -61.87 | 39.22 | -63.88 |
| 14 Life Below Water | -2.50 | -40.59 | -79.19 | -60.82 | 33.14 | -123.33 | -35.46 | 27.05 | -49.46 |
| 15 Life on Land | -2.45 | -55.71 | -101.54 | -104.65 | 33.06 | -154.17 | -58.73 | 30.97 | -66.23 |
| 16 Peace and Justice Strong Institutions | -7.65 | -76.87 | -137.98 | -260.00 | 40.16 | -235.14 | -61.24 | 39.61 | -97.42 |

Source: Web of Science.
Colour code of the Table reflects relative performance on SDG-related metrics for Brazilian PhDs living in Brazil (blue) vs. abroad (pink); deeper colours indicate higher performance.
% Docs Cited: Share of published papers; CNCI: Category Normalized Citation Index; % Top 10%: Papers in the top 10% by citations; % Ind. Collab: Industry collaborative papers; % Corresp. Author: Papers with PhD as the corresponding author; % Highly Cited: Papers frequently cited in a field; IRW: World relative impact; % Gold Docs: Papers in Gold Open Access; % Q1: Papers in top-quartile journals.

## Discussion

Publishing is a basic science and research activity (Blind et al. 2018). This approach has already been used in the statistical accounting of scientific migration flows: recently, the OECD journal Science, Technology, and Industry Scoreboard (OECD, 2017) has been tracking the movement of scientific personnel at the national level using Scopus data. Moed et al. (2013) used the methodology to analyze the migratory trajectories of scientists from the United States, Germany, United Kingdom, Italy and the Netherlands. The bibliometric approach has yet spread so far. Still, it has the significant potential both for collecting the contacts of representatives of a diaspora and for statistically monitoring the entry and exit of scientific personnel.

A large number of scholarships abroad, but without appearing in the list of publications, doctorates and orientations, refers, in large part, to fellows in the Science without Borders program, which operated between the end of 2011 and 2016, when the last fellow went abroad (McManus & Nobre, 2017). Koslowski (2014) identified the USA, Australia, Canada and the United Kingdom as the most prominent talent attraction countries, in line with the results here and the leading countries where Brazilian authors maintain international collaboration (McManus et al., 2020). The importance of the training year after 2012 may be due to the increase in scholarships abroad after this period because of the Science without Borders program (McManus & Nobre, 2017), and the fact that the REUNI program (Programa de Apoio a Planos de Restructuring and Expansion of Federal Universities) which began in 2008 and hired new doctors after this date (Paula et al., 2020). In recent years, there has been a reduction in opportunities for hiring new doctors. Still, the postgraduate system continues to expand with the formation of more than 20,000 new doctors each year (McManus et al., 2023), thereby increasing competition and creating the need to look for jobs elsewhere.

The important Brazilian Universities identified align with Brazilian universities’ performance in international rankings (Prado, 2021). Internationalization is a requirement for postgraduate courses with the highest scores (6 and 7) in Brazil in the Capes quadrennial assessment (Paiva and Brito 2019), exposing a more significant number of its students to international experiences. Factors leading to stabilising the number of Brazilian diasporas may include Brexit, where immigration limitations for postdocs in the UK increased competition in other countries (Kratenova, 2019), the Covid-19 pandemic, and the reduction of scholarships from Brazil to study abroad.

Brazil can benefit from this knowledge produced by the diaspora abroad. For example, The AI Index Report 2023 (Maslej et al., 2023) shows that although Brazil elaborated a national strategy in 2021, the country has no significant system and no significant system author (p. 52-53). Yang and Welch (2010) coined the term knowledge diasporas to explain highly skilled mobility, underpinned by increased global migration flows and information and communication technologies’ rise (ubiquity). This knowledge diaspora has become an essential channel for technology transfer from the host country, mitigating the adverse effects of brain drain in the home country (Welch and Zhen, 2008). When examining US patent data, Breschi et al. (2017) conclude that this diaspora effect is significant among migrants born in Asia. However, the brain gain effect (inventors residing in the home country) is insignificant for India. The transnational diaspora of the highly skilled can benefit the global knowledge base by narrowing the North-South scientific and technological gap, as evidenced by Chinese knowledge diasporas (Zweig, 2006). Themes such as machine learning and covid-19 were important in the present study and are recognized worldwide with the onset of the pandemic in 2020 (Akomea-Frimpong et al., 2022; Burkov, 2019).

Transformations occur in the inclinations and habits of individuals and also occur in the priority of production, resulting in the generation of knowledge and companies of a new character, which is at the forefront of the economic scenario, such as the technical sector and service sectors such as Microsoft, Apple, Amazon, Alphabet, Facebook, Alibaba. In addition, giant companies retreated in the automotive, oil and gas sectors. As a result of this transformation, physical capital has shrunk dramatically and has been replaced by human capital. Knowledge, creativity and innovation constantly evolve because they are related to the human element, which increases with experience over time, thanks to education and training (Braunerhjelm, 2010). Therefore, Brazil needs to think about recapturing the knowledge it invested in with the diaspora.

The return migration of highly skilled personnel in areas of importance for the country’s development (Table 5), especially from the Global North to the Global South, has attracted increasing interest from policymakers and researchers (Zweig and Wang, 2013). For example, Choudhury (2015) confirmed that migrants returning to the Indian R&D hub of multinational companies helped promote technology diffusion and facilitate activities. As a global innovation ecosystem emerges, the emerging diaspora of STEM talent networks is potentially a game-changing phenomenon that affects where, by whom and how innovation activities are performed (Lewin & Zhong, 2013). Skilled migration will likely continue to be encouraged by OECD countries (Alleyne & Solan, 2019). Developing countries, such as those in Latin America, including Brazil, should seek to benefit from this migration.

Machine Learning techniques were developed to analyze high-throughput data to gain actionable insights, categorize, predict and make evidence-based decisions in new ways, which will drive the growth of new applications and fuel the sustainable growth of AI (Xu et al., 2021). The Government Artificial Intelligence Readiness Index 2019 (Oxford Index) shows that Brazil has an index of 2/3 of most OECD countries; therefore, the diaspora could be used strategically.

França & Padilla (2018) and Souza & Iorio (2018) highlight that the resurgence of Brazilian emigration in Portugal has differentiated characteristics with numerical expressiveness but a diversity of profiles. This new migratory flow involves a new profile of Brazilian immigrants, composed of individuals with greater financial and entrepreneurial capacity, a high level of professional qualification or individuals seeking higher qualifications (master’s students, doctoral students and researchers). This preference may be due to the common mother tongue (Lipski, 2022) and the ease of obtaining nationality due to historical roots (IBGE, 2023).

One factor that affects the intensity of scientific migration and knowledge transfer, particularly important, is the sharing of cultural traits, mainly a common language (Yurevich et al., 2019; MacGarvie, 2005). In this sense, the frequency of diaspora in countries such as Portugal, Germany and Italy would be expected. There is the enduring significance of national history and culture, and difficulties associated with establishing an academic career internationally. However, it is crucial to consider the large number of academics who are working abroad. Data from the recent “GlobiSci” survey of more than 47,000 academics working in biology, chemistry, earth and materials sciences in 16 countries provide a snapshot of international academic work (Franzoni et al. 2012). The results show that 36% of the sample worked outside their country of origin. An even larger share of academic scientists in Switzerland, Australia and Canada are foreign nationals. Still, the US attracts the largest number of scientists from abroad and is likely among the biggest beneficiaries of net academic inflows. The reasons that led them to take up a position outside their country of origin revealed the most popular answer as the “opportunity to improve future career prospects” (Franzoni et al. 2012). This suggests that most internationally mobile scholars are in the early stages of their careers. The response supports a push-pull account, which claims that lack of opportunity at home drives academics to seek opportunities abroad. Such an account is also consistent with what Ackers (2008) has described as “forced mobility”, in which young academics from resource-poor countries must move to pursue an academic career. These interpretations can lead to the conclusion that mobile academics depend on their hosts.

The interconnection between migrations and political and bureaucratic factors remains a debatable issue. For example, a recent study based on an analysis of the visa legislation of 38 countries for 1973-2012 demonstrates that visa limitations in recipient countries substantially reduce the number of short-term visits and, at the same time, cause people to move to these countries for an extended period or forever. However, while the introduction of visa restrictions affects international migration with a considerable delay (migration flows decrease by 20% in 10 years, on average), the consequences of visa abolition manifest themselves significantly more quickly (on average, the number of migrants increases by 30% in three years) (yang & Welch, 2010). According to some studies, visa limitations negatively influence not only the intensity of migration of scientists between countries but also the number of international scientific collaborations (Appelt et al., 2015).

Brazilian science funding is dispersed, with resources from 68 countries and 696 funding sources being cited (McManus & Baeta Neves, 2021). Ten countries, the European Union and 80 companies were responsible for 90% of the articles produced by Brazilian researchers. The USA and Germany are the main foreign funders of Brazilian documents, with 6.8 and 2.3%, respectively. Accounting for all European collaboration, it surpasses US collaboration (50% more).

One might work as a postdoc for several reasons, including an abundance of doctoral-level researchers concerning permanent academic positions (Stephan 2012) and rising expectations in some fields that new researchers spend at least some time as a postdoc (Goldman and Marshall 2002). Evidence shows that researchers working as postdocs tend to be more productive in research than their peers without postdoctoral experience (Horta 2009; Su 2011). Given this, some academics may choose to work as postdoctoral fellows to improve future employment prospects. Roos et al. (2014) indicated that researchers who did a full doctorate in Brazil published more articles than those abroad. However, articles by researchers who completed their doctorates in Brazil were published in journals with a lower impact factor. They received fewer citations than articles published by researchers who completed their doctorates abroad. The results indicate that the qualitative performance of researchers who did their doctorate abroad was better than those who did their doctorate in Brazil. Consequently, Brazilian government policies must be geared towards increasing the relevance of Brazilian articles regarding scientific quality and international insertion.

Qualitative research on international postdoctoral employment in the US and UK supports the notion that universities employ international postdocs because they are productive in research. It also provides insight into hiring considerations (Cantwell 2011a, b). In Brazil, this experience may not be considered since the competition system limits the participation of advisors and supervisors in selecting candidates, and hiring is usually to offer undergraduate classes rather than research and graduate potential. Postdoctoral fellows are among the fastest-growing group of academic staff in the United States, and more than 50% of all postdocs in the United States are temporary visa holders (Cantwell & Taylor, 2013).

Studies show that female scientists with experience abroad came back with new identities shaped by their life and education abroad and by their university exposure to people from different cultural backgrounds (Alkubaidi & Alzhrani, 2020). They also got used to a more comfortable lifestyle in their host countries.

A recent study assessing internationalization among academic scientists found that 38% of samples in the United States (US) came from abroad (Franzoni et al. 2012). Early career scholars are especially mobile. Every year, many internationally mobile scholars come to the US to work as postdoctoral fellows. In 2009, the majority (53%) of the approximately 60,000 postdocs employed at US universities were foreign nationals (National Science Foundation [NSF] 2012). This means that postdocs are the most internationalized group in US higher education. The national-level conditions highlighted by the “push-pull” models are not the only determinants of international academic mobility. Global and local dynamics, individual and institutional circumstances, and experiences also shape the fields in which actors operate (Marginson 2011; Marginson and Rhoades 2002). Research on the employment of international postdocs at universities in the United States and the United Kingdom (UK) found that individual personal and professional circumstances, institutional policy, and material job titles influenced the hiring process (Cantwell 2011a). In addition, early career scholars and principal investigators (“PIs”) believe that international employment demonstrates world-class competitiveness. This shared vision shapes the motivations of internationally mobile postdocs and their employers (Cantwell 2011b).

Globalization has led to increased academic mobility across national borders. International academic mobility is often conceptualized using a “push-pull” framework in which local deficits “push” individuals away from their home country and desirable conditions “pull” them to specific locations abroad (Altbach 2004; Mazzarol and Soutar 2002). Such accounts postulate that the lack of resources and opportunities in one context relative to another explains the movement of scholars across national borders. While this model helps describe aggregate flows, it has limitations in fully explaining academic mobility (Lee and Rice 2007; Lee et al. 2006). For example, “push-pull” models do not consider the “demand” in host countries for internationally mobile scholars.

## Concluding Remarks

Retaining, engaging and recruiting qualified professionals from their diaspora are not the object of policies exclusive to countries in the Global South. The Canadian Institute for Advanced Research, established in 1982 with an annual budget of $25 million, aims to retain Canada’s best and brightest researchers by supporting research programs and funding more than 400 researchers. The International German Academic Network aims to attract German scientists in North America by offering job placement assistance and research funding. German academic organizations have funded up to €100,000 to German universities to help them attract and hire German academics (Cohen and Kranz, 2015). However, diaspora recruitment policies are not more widely used by the Global South (Hooper and Sumption, 2016) than those of the Global North, even considering that, in some cases, they represent the only political alternative to attract highly skilled immigrants.

A diaspora network can be a source of new dynamism for the economic development of any country (e.g. Canada and Australia, whose immigration policies favor immigrants with advanced STEM training) and has recently become the focus of national policy in emerging economies due to the potential that STEM diasporas have for the economic development of their country of origin. For example, Indian emigrants in the US are mostly highly educated and many of them are specialists or managers in high technology companies (Pew Research Center, 2012). Financing agencies in Brazil must take this into account when building their internationalization policies. One of its concepts is that the design, implementation, monitoring and evaluation of migration policies must be considered in the broader context of relevant sectoral development policies. The interrelationships strongly depend on the country context and the conditions of implementation of different programs, and are reflected in migration policies and results in the recruitment of highly qualified people from the diaspora (li et al., 2018). While two-thirds of OECD countries had policies to attract highly skilled migrants in 2015 (Czaika and Parsons, 2017), a 2011 global survey found that fifteen of the world’s top twenty-five migrant source countries have policies that encourage migration of return (Hooper and Sumption, 2016).

As Brazil in general invests less in STEM (Science, Technology, Engineering, Mathematics) or STEAM (Science, Technology, Engineering, Arts, Mathematics) and the knowledge area needs more resources to carry out highly complex projects, access to funding and career options is smaller for researchers (Valenzuela-Toro and Viglino, 2021) The migration of scientists and STEAM higher education students to high-income countries is therefore a significant challenge affecting human capital formation. This phenomenon, commonly known as brain drain, results in the loss of some highly trained or skilled individuals. The social impact of displacement not only reduced the economic potential of the region (Cerovic and Beaton, 2017), but also reduced technological innovation (Lozano-Ascencio and Gandini, 2012). is reflected in the low production of patents and publications (Ciocca and Delgado, 2017) and the decrease in access to high-quality education (Busso and Messina, 2020).

Mahroum (2000) observes that, for academics and scientists, scientific faculties and norms operate a self-organized movement between and between research institutions, therefore, the reputation of scientific openness, excellent quality and prestige for excellence in research have been shown to be important qualities. to all organizations seeking to attract high caliber scientists; while engineers and technicians are driven primarily by economic factors and will go where the demand for their skills is most needed and best rewarded. In the case of Brazil, it is shown that there are hiring norms in public universities (where there is more than 90% of research in Brazil) that rule against research productivity since professors enter at the lowest level regardless of the quality of the curriculum or previous experience, there is equality of wages within the country, which leads fewer and fewer new teachers to participate in research (Baeta Neves et al., 2020). Research grant values are also not attractive (McManus et al., 2023). In agreement with Marmolejo-Leyva et al. (2015), mobility has a strong impact on international scientific collaboration. We found no substantial influence among researchers who obtained their doctoral degrees abroad and those who graduated from Brazilian universities.

To build more meaningful collaborations with partners in Brazil, to be successful these efforts not only require strong institutional support from departments and universities in Brazil and abroad (which Brazil appears to lack), but also depend on the erasure of long-standing prejudices. date. Without this, the collaborative landscape can be a difficult, exhausting, and lonely endeavor, whose success and impact is limited to what individuals can achieve. Furthermore, there are significant career implications of single-handedly coordinating long-term collaborative processes that include high research transaction costs, slower and lower publication rates, and unsustainable work-life balance (Sellberg et al., 2021). These setbacks are universal and well-articulated in a growing body of literature on the challenges of doing transdisciplinary research in traditional university settings (Moore et al., 2014; Patterson et al., 2013; Sellberg et al., 2021).

Despite continuing asymmetrical power relations between the global North and South, the global race for talent is no longer a privilege enjoyed only by developed countries. Comparative analysis of the content and effectiveness of policies has attracted increasing interest. Examination of the policy initiatives that result in the return migration of highly skilled migrants from the Global North to the Global South, however, remains inadequate. Several points raised here by Brazilians abroad are in agreement with the problems found in HEIs in Russia by Oleksiyenko (2021).

The experience of India and especially of Iran shows that, even in the absence of a consistent State policy regarding the scientific diaspora, emigrant scientists are actively helping to develop the scientific and technological potential of their homeland, maintaining contacts with colleagues, visiting their country of origin give lectures and participate in scientific collaborations. Associations of fellow scientists, which are often formed without the direct participation of the State, play an important role in this process. In this situation, it remains for the State to establish contacts with them and create comfortable conditions for their interaction with scientists and scientific organizations at the national level. Existing programs for recruiting foreign researchers to work on temporary projects are aimed at representatives of these associations. At the same time, the experience of Korea, Taiwan and China shows that the mass remigration of highly qualified scientists and specialists requires somewhat different mechanisms and direct state participation. Measures to materially encourage repatriation proved to be the most effective: competitive salaries for top scientists and engineers; the creation of science parks and special economic zones, where specialists are invited to work under attractive conditions (for example, in some Chinese regions, returning specialists enjoy benefits when receiving a loan or buying housing); and support for businesses focused on innovation, which, in turn, invest in R&D and hire specialists from abroad.

The opinions expressed by Brazilians abroad are in line with those of other countries (Alkubaidi and Alzhrani, 2020), where experience in other countries is not welcome in Brazil. According to the OECD (Auriol, 2010), the labor market for doctoral graduates is more internationalized than that of other tertiary-level graduates, and the doctoral population is highly internationally mobile. In European countries for which data are available, 15% to 30% of PhD holders who are citizens of the country reported experiencing mobility abroad during the last ten years. Migration and mobility patterns of doctoral students are similar to those of other higher education population categories and others with important flows to the United States, mainly from Asian countries, and large intra-European flows, notably to France, Germany and the United States Kingdom.

The present study has recognized limitations when (i) it does not identify former students not publishing abroad, such as, for example, professionals such as doctors, engineers, etc. (Frehywot et al., 2019) and (ii) it does not identify people with name changes (e.g. social name or by marriage).

## Material and methods

Information was collected on students completing a doctorate in Brazil between 2005 and 2021 from Capes open data web page and the names of scholarship holders. Information was also available on those with a full doctorate abroad or postdoctoral scholarships on the Capes, CNPq and FAPESP websites.

We searched for diaspora in two manners:

i. The names of former students with only Brazilian affiliations were deleted, and the rest were searched in Scopus® (Elsevier) to verify affiliations abroad. The researchers’ information was downloaded in CSV format, and the files were joined in Excel and analyzed in Vosviewer®.
ii. In Incites (based on Web of Science), the names of researchers affiliated with Brazil between 2005 and 2022 were collected. Afterwards, researchers with only Brazilian affiliations were deleted. Once the number of researchers exceeded the download limit (50000) the download was carried out in groups. Then the files were combined in Excel, and Web of Science ReseacherID and ORCID used to eliminate duplicates.

In this period, 452,595 Brazilian researchers published 748,374 documents. Postgraduate professors/supervisors (Table 1) were eliminated from both lists as it was presumed to be in this category they reside in Brazil (this information is also available in Capes open data).

The methodology used in this study is not new. Bibliometric information provides rich possibilities for considering the circulation of scientific personnel on a global scale of science: a scientific publication contains information about the author’s affiliation, a change to a foreign affiliation implies the scientist’s departure from the country. According to the methodology (OECD, 2017; Moed et al., 2013) an author was considered representative of the diaspora if he had at least two affiliations within the specified period. At least one of them had a relationship with a Brazilian scientific organization (a research institute or a higher education establishment), and the other with a foreign one.

Groups of PhDs that graduated with Brazilian funding between 2005 and 2021 were formed:

- All PhDs graduated in Brazil.
- PhDs with a sandwich period abroad
- PhDs paid to do a full PhD abroad
- Postdoctoral experience abroad.

Those without affiliation in Brazil were then identified in Scopus and Web of Science. Data were visualized in Vosviewer®. Afterwards, they were analyzed with logistic regression and decision tree (0 – without address abroad and 1 – with address abroad) in SAS (Statistical Analysis System Institute, Cary, North Carolina) to verify the effect of year of graduation, area of training (Social and Human, Life or Exact sciences), Legal Status of the HEI (Federal, State, Municipal or Private), and post-graduate program score. We also tested the effect of TOEFL score (where available) but this was not relevant and so we removed it from the analysis.

## Author contributions

C.M., B.A.D.N., A.A.B.N., R.T.S., and C.P.F., and conceived the study and contributed to study design and analyzed the data. C.M., B.A.D.N., and C.P.F. wrote the manuscript. All authors read and approved the final version.

## Data Availability Statement

The datasets generated and/or analyzed during this study are available in Scopus and Web of Science; however, access to these datasets may be restricted as they are behind paywalls. Additional data supporting the findings of this study can be obtained from the corresponding author, Concepta McManus, upon reasonable request.

## Competing interests

The authors declare no competing interests.

## Footnotes

2 https://www.oxfordinsights.com/ai-readiness2019

## Notes

### Competing Interest Statement

The authors have declared no competing interest.

